# Double UP: efficient *in utero* electroporation via internal controls

**DOI:** 10.1101/571323

**Authors:** Russell J. Taylor, Justin Carrington, Kendra L. Taylor, Karl E. Richters, Leah R. Gerlach, Erik W. Dent

## Abstract

*In Utero* Electroporation is a powerful tool for testing the role of genes in neuronal migration, but this technique suffers from high degrees of variability due to multiple factors. Therefore, we developed Double UP, a novel approach that combines LoxP-flanked reporters and limiting Cre dosages to generate internal controls. This technique allows for more rigorous quantification of migration, while decreasing the number of animals, reagents and time to complete experiments.

---

The method of *In Utero* Electroporation (IUE) ^1, 2^ has contributed greatly to our understanding of neuronal differentiation and migration in the brain. Indeed, it is the *de facto* technique to determine how neuronal function or migration is disrupted in the central nervous system after genetic manipulation or in disease models. However, this technique suffers from a high degree of variability due to inherent challenges, including inconsistent surgeries, developmental differences within and between litters and gradients of maturation within the developing brain ^3^. These sources of variability are cumulative, leading to difficult to interpret data and an increased risk of both false positives and false negatives. Furthermore, postnatal analysis of brains is difficult as researchers must either abandon littermate controls entirely, or use additional plasmids to indicate which animals were control or experimental.

To address and mitigate the variability inherent to IUE, we have developed a dual-fluorescent plasmid, designed to generate an internal control. This plasmid, termed “Double UP”, uses the strong ubiquitous CAG (<u>C</u>MV enhancer/beta-<u>A</u>ctin promoter and rabbit <u>G</u>lobin PolyA tail) promoter ^4^. The CAG promoter is followed by a LoxP-flanked cassette ^5^ consisting of the green fluorescent protein mNeon-Green ^6^, a stop codon and the rabbit-globin PolyA tail ^7^. Following the second LoxP site is a red fluorescent protein (mScarlet) ^8^, with identical stop codon and rabbit-globin PolyA tail **(Fig. 1a)**. In the absence of Cre, cells containing this plasmid produce mNeon-Green **(Fig. 1b)**. However, if Cre is present in the cell, the mNeon-Green is excised, and replaced in the exact same locus by mScarlet, with an identical polyA tail. Co-injection with a limiting dose of pCAG-Cre ^9^ plasmid allows rough control of the ratio of green cells to red cells. Very low amounts of Cre plasmid result in a few scattered red cells, moderate amounts of Cre plasmid result in roughly half red cells and half green cells **(Fig. 1b)**, and high doses of Cre plasmid result in all red cells. A molar ratio of approximately one copy of Cre plasmid for every 75 copies of Double UP results in roughly 50% green and 50% red cells. Thus, approximately equal numbers of red and green cells can be labeled in the same cortical section.

**Figure 1:**
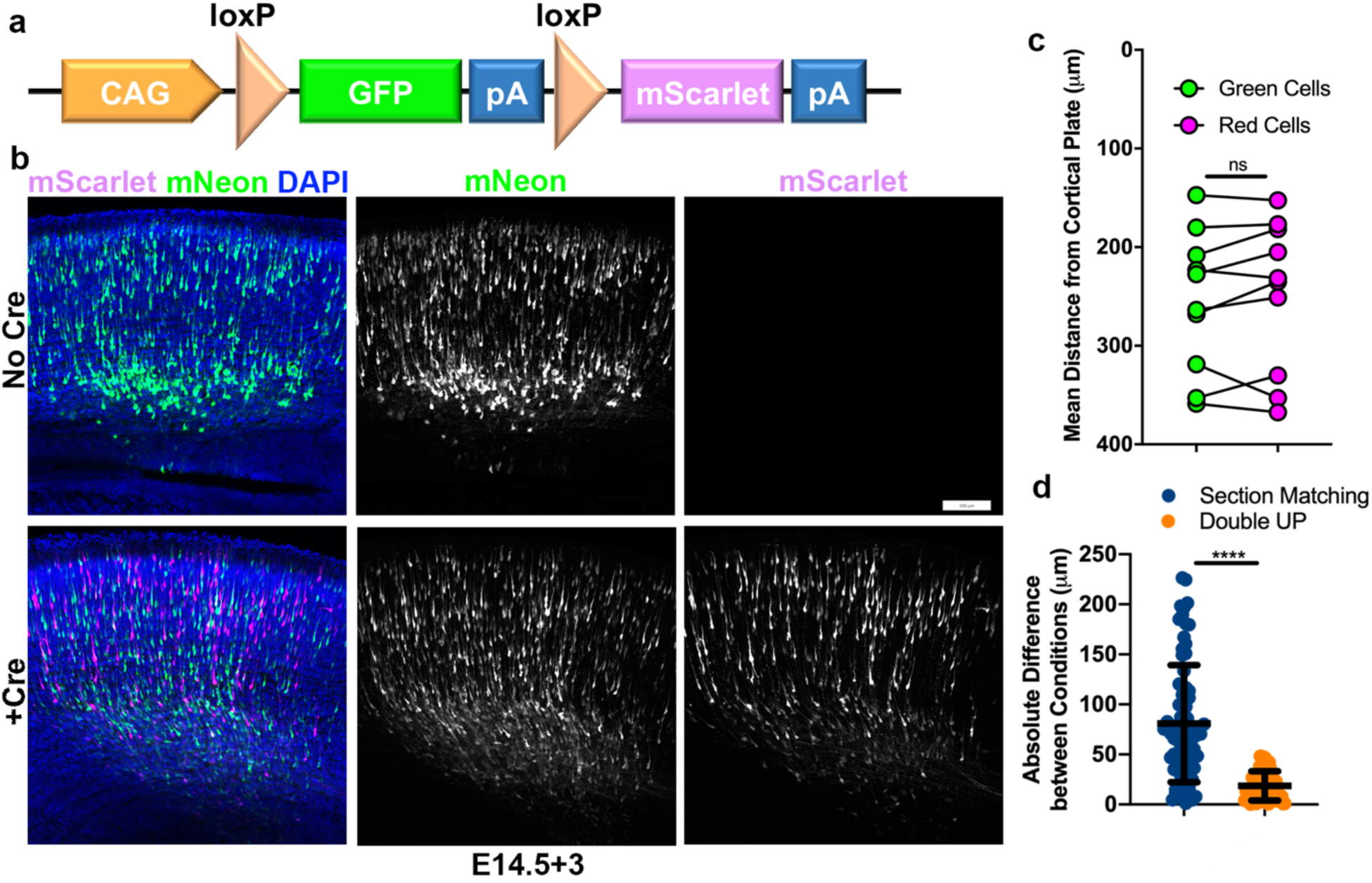
Construction and characterization of Double UP. **(a)** Schematic of Double UP construct. **(b)** Representative IUE of Double UP without Cre (top) and with pCAG-Cre in a 75:1 molar ratio (bottom). Embryos were electroporated at E14.5 and allowed 3 days to mature (E14.5+3). Scale bar, 100µm. **(c)** Comparison of migration of mNeon-Green- and mScarlet-positive neurons, E14.5+3. Each dot represents mean distance from the top of the cortical plate of neurons within one slice. Connected dots indicate that those measures came from the same brain. n=10 slices with 100 - 976 neurons from each slice. No significant difference between green and red populations (two-way ANOVA). Error bars not shown for clarity. **(d)** Comparison of reliability of controls between section matching and Double UP. Full data in Supplementary Figure 2. 76 comparisons made for section matching, 61 for Double UP. Only sections that had a perfect match in another brain were included for either analysis. **** p<0.0001., Kolmogorov-Smirnov t-test, two-tailed. Lines indicate mean and SD.

We wanted to rigorously determine if the mNeon-Green and mScarlet fluorescent proteins affected neuronal migration relative to one another. This was done to demonstrate the validity of using green as an internal control to red, by demonstrating that in the absence of manipulation the green and red cells behave similarly. To produce more quantitative and consistent data than is currently used in most neuronal migration studies, we developed a program which allows for automated quantification of cell body location and brightness, as well as distance from the nearest point on any user-defined region of interest. This Java-based program, entitled “TRacking Overlapping Neurons”, or “TRON” utilizes simple processing steps, followed by a modified version of the Fiji plugin 3D Object Counter. TRON identifies the center of mass of every fluorescently-labeled neuronal cell body and calculates the distance to a user defined region of interest (ROI) (**Supplementary Fig. 1a-g**). These data can be further split into bins (**Supplementary Fig h-j)**. The individual processing steps are available in Online Methods, and the TRON program and associated instructions are downloadable at https://uwmadison.box.com/v/tron. Implementing this program, we were able to determine that mNeon-Green and mScarlet-labeled neurons migrate in statistically similar fashions in embryonic brain **(Fig. 1c)**. Because all data from Double UP consists of matching sets of green and red cells, more powerful statistical tests (two-way ANOVA) can be used to determine if red and green neurons are behaving differently. Together, these data indicate that Double UP is a viable technique to use to compare control and experimental conditions in single brain sections.

One confounding variable in traditional IUE is the requirement of using matching sections in control and experimental brains. It is well established that embryonic cortical development proceeds along both rostral-caudal and medial-lateral gradients ^3^. However, the degree to which these gradients cause variabilities to IUE has not been well documented. To determine this, we performed IUE at E14.5 and collected brains three days later, in the midst of migration. We imaged and quantified migration for thousands of cells in every sequential 100µM coronal section of brain (n=61 sections, 4 embryos, from 2 pregnant mice). Distance was measured from top of cortical plate for each cell, as a measure of distance from completion of migration **(Fig. 1d, Full data Supplemental Figure 2)**. Migration data were analyzed using either intra-sectional variation using Double UP, or inter-sectional variation using matched sections. There was more than four-fold more variation comparing section-matched sections (mean of 80.8+/−6.7μm (SEM)) than within sections using Double UP (mean of 18.6+/−1.9μm (SEM)). If even neighboring sections, or single sections, demonstrate high degrees of variability, it calls into question the practice of using matching sections from separate brains as controls.

**Figure 2:**
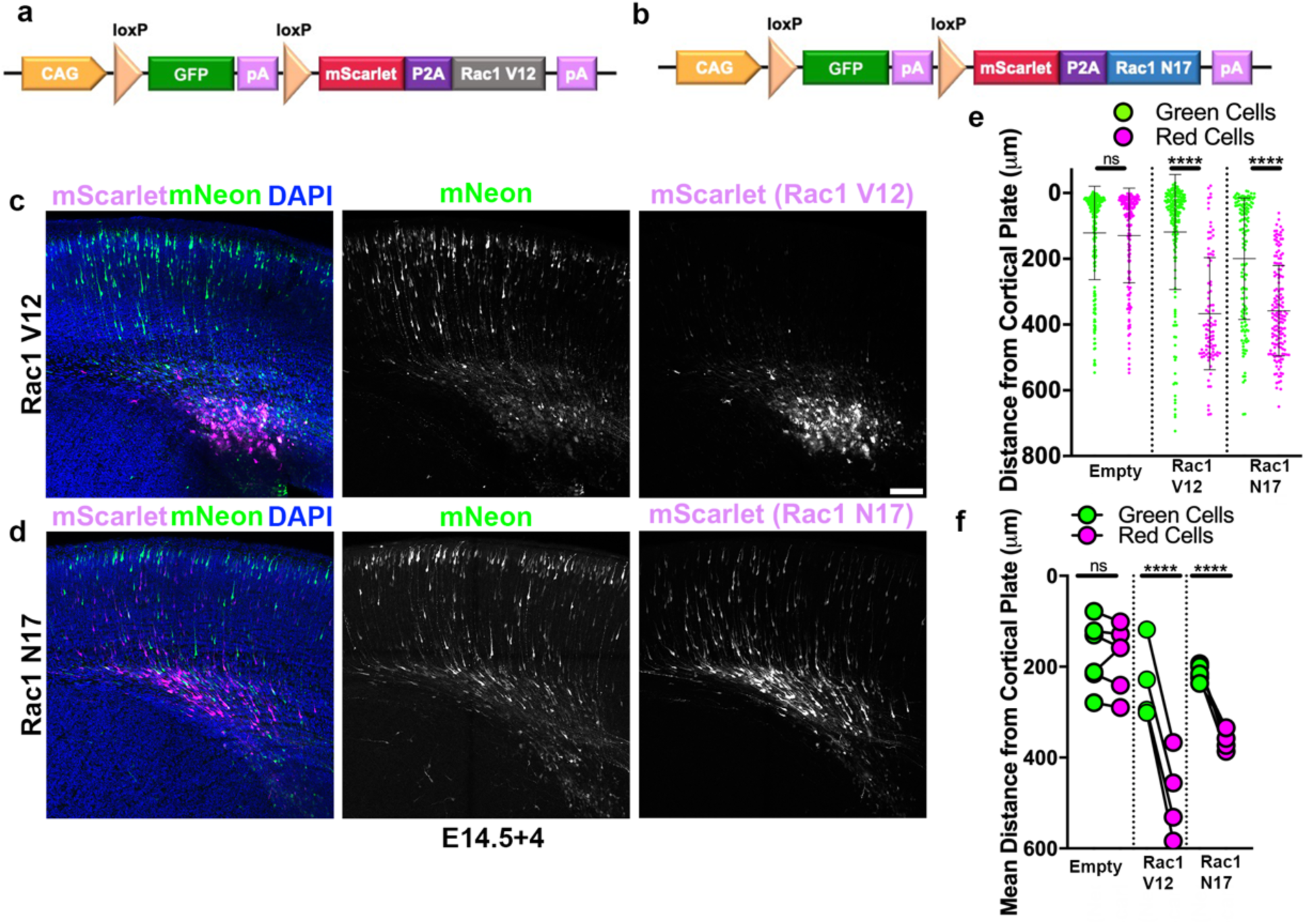
Double UP Replicates previous findings. **(a)** Schematic of Double UP Rac1-V12. **(b)** Schematic of Double UP Rac1-N17. **(c)** Representative image of Double UP Rac1-V12, E14.5+4. Scale bar, 100µm. **(d)** Representative image of Double UP Rac1-N17 E14.5+4. **(e)** Dot plot of three representative slices from Double UP Empty, Double UP Rac1-V12 and Double UP Rac1-N17. Each dot represents distance from the top of the cortical plate to the center of mass for each neuron. ns=not significant, **** p<0.0001, Kolmogarov-Smirnov t-test, two-tailed. Lines indicate mean and SD. **(f)** Dot plot of Double UP Empty, Double UP Rac1-V12 and Double UP Rac1-N17 (n=6 slices, 4 slices, 5 slices respectively). Each dot represents mean distance from the top of the cortical plate to the distance of all cortical neurons in a slice. Connected dots indicate measurements were made in the same brain. ns=not significant, **** p<0.0001, two-way ANOVA, using Two-stage setup of Benjamini, Krieger and Yekutieli. Error bars not shown for clarity.

The intent of Double UP is to introduce manipulations in combination with the red fluorophore to distinguish if and how the red experimental cells differ from the green control cells. To accomplish this, it is vitally important that Cre-negative (green) cells do not express any red fluorescent protein or any manipulation. It was also important to test the system for “leakiness” using ERT2-Cre-ERT2 ^9^ (pCAG-CreER) in the absence of tamoxifen, to inform future experiments with delayed activation of Double UP. To test for leakiness, we undertook multiple tests. First, IUE was performed with Double UP and either no Cre, pCAG-CreER (75:1 molar ratio) or pCAG-Cre (75:1 molar ratio). Imaging settings were determined using the pCAG-Cre electroporations, and unchanged between sections. Rather than using automatic thresholding, images for this set of experiments were artificially thresholded at 1000 gray values. In the absence of Cre, there was a single weakly positive cell across three sections from three different brains. IUE of Double UP and pCAG-CreER resulted in 3-4 positive cells in each section, and IUE of Double UP and pCAG-Cre resulted in roughly equal populations of green and red cells **(Supplementary Fig. 3a)**.

However, absence of individual bright cells does not prove Double UP is not leaky, so we next quantified the intensity of red signal of individual fluorescent cells. Fluorescent intensity measures were collected for 90 green cells in no-Cre and CreER conditions, and from 90 red cells in a +Cre condition, from three different brains per condition. Equivalent measures were also taken for equivalent cells in the contralateral cortex of the no-Cre and CreER brains, which did not receive plasmid. Surprisingly, red values were subtly but significantly lower in the electroporated side of no-Cre cortex, compared to the contralateral side **(Supplementary Fig. 3b)**. The reason for this difference is not clear, but importantly the values were not higher than the electroporated side of the cortex. Finally, we performed Western blotting for detection of any protein associated with mScarlet. To accomplish this, we used Double UP 3x-HA, in which 3x-HA was fused with the mScarlet fluorophore **(Supplementary Fig. 3c)**. Western blotting analyses performed in a mouse catecholaminergic cell line (CAD cells), demonstrated that in the presence of very little Cre plasmid, HA protein is readily detectable, but in the absence of Cre there is absolutely no detectable HA protein **(Supplementary Fig. 3d, e)**. Together, these data suggest that in the absence of Cre, no mScarlet or associated protein is produced.

In the absence of manipulation, red and green neurons migrate equivalently **(Figure 1C)**. This allows for attaching a manipulation of interest to the red fluorophore and examining the effects of this manipulation relative to the unmanipulated green neurons. Thus, we set out to determine if using Double UP recapitulated previously published findings from multiple groups. Both constitutively active Rac1 (Rac1-V12) and dominant negative Rac1 (Rac1 N17) have been previously shown to inhibit neuronal migration ^10, 11^. Rac1-V12 and Rac1-N17 were separately cloned downstream of mScarlet, following a P2A peptide **(Fig. 2a, b)**. The P2A causes a ribosomal skip during translation, so that equimolar ratios of both mScarlet and either Rac1-V12 or Rac1-N17 are generated. Cells receiving both Double UP and Cre plasmids now produce mScarlet and also untagged Rac1-V12 or Rac1-N17. Cells receiving only Double UP will produce only mNeon. Following IUE of Double UP Rac1-V12 or Rac1-N17, red neurons fail to migrate, while green neurons migrate normally **(Fig. 2c-f)**. Consistent with previous findings, while both gave similar migration profiles, Rac1-N17 neurons much more closely resembled the morphology of adjacent green neurons than did the Rac1-V12 neurons (data not shown).

*In Utero* Electroporation is powerful and widely used but is traditionally dependent upon extremely accurate and insufficient section matching. Brains have strong developmental gradients acting along multiple axes, increasing the difficulty of accurate section matching with regards to traditional IUE. Experiments can now be performed within a single embryo, from a single surgery. Animal to animal variation is no longer an issue, relatively minor inconsistencies in surgical technique are made inconsequential and section matching is obviated. Implementation of Double UP can provide more rigorous data, while simultaneously reducing costs and animal numbers.

## ACKNOWLEDGEMENTS

We thank all the members of the Dent lab for helpful discussions and critical comments on the manuscript.

pCAG-Cre was a gift from Connie Cepko (Addgene plasmid # 13775; http://n2t.net/addgene:13775; RRID:Addgene_13775). pCAG-ERT2CreERT2 was a gift from Connie Cepko (Addgene plasmid # 13777; http://n2t.net/addgene:13777; RRID:Addgene_13777). The project was supported by NIH grants RO1-NS080928 to W.W.D., T32-GM007507 to K.L.T and R.J.T. and by the Clinical and Translational Science Award (CTSA) program, through the NIH National Center for Advancing Translational Sciences (NCATS), grant UL1TR002373. The content is solely the responsibility of the authors and does not necessarily represent the official views of the NIH.

## AUTHOR CONTRIBUTIONS

R.J.T and E.W.D. conceived the project. R.J.T., J.C., K.L.T., L.R.G. executed experiments. R.J.T and E.W.D. wrote the paper with input from all authors. E.W.D. supervised all aspects of the work.

---

**Supplementary Figure 1:**
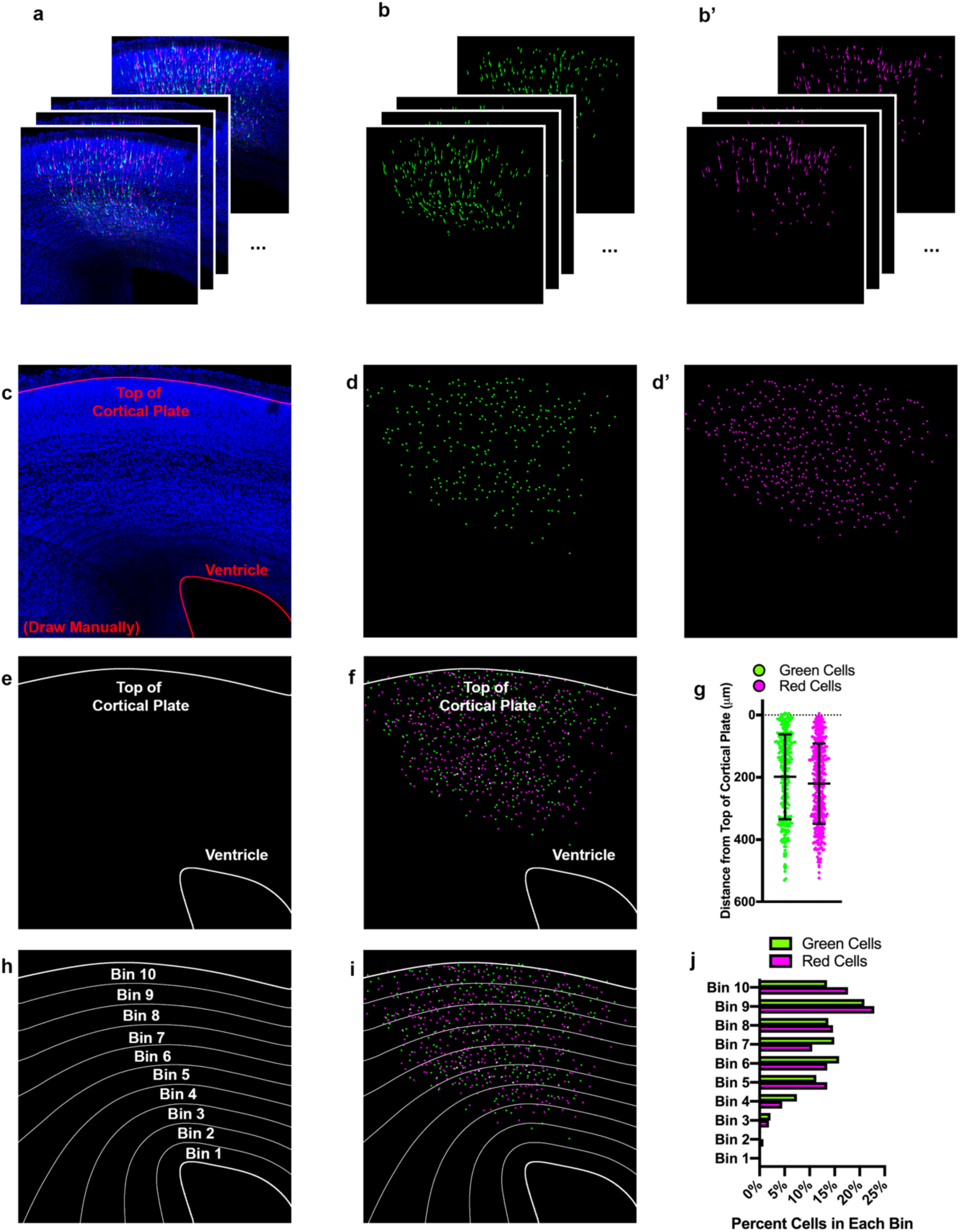
Automated tracking of cellular location with TRON program. **(a)** Example input. 12, 3-color confocal slices. **(b)** Green and red colors, processed independently and automatically via: unsharp mask, gaussian blur, conversion to 8-bit, automatic threshold, erode, watershed. **(c)** Manual drawing of regions of interest (ROIs – red lines at top of cortical plate and ventricle). This is the only manual step in the process. **(d, d’)** Output of modified 3D Object Counter plugin from Fiji. Center of mass of each cell is identified and mapped; X, Y location and brightness of original is obtained and recorded. **(e)** Location of each cell mapped onto the ROIs. **(f)** Distance from the center of each cell to the nearest spot on each ROI is measured and recorded. **(g)** Graphical representation of distance each cell is from top of the cortical plate. Cells located above the top of the cortical plate are treated as having a “negative” distance. Lines indicate mean and SD. **(h)** Distance between ROIs can be split into bins, calculated to reflect the ROI to which they are closest. **(i)** Location of each cell mapped onto the bins. **(j)** Graphical representation of the data split into 10 equal-sized bins, each bar shows percentage of cells of that color in the respective bin.

**Supplementary Figure 2:**
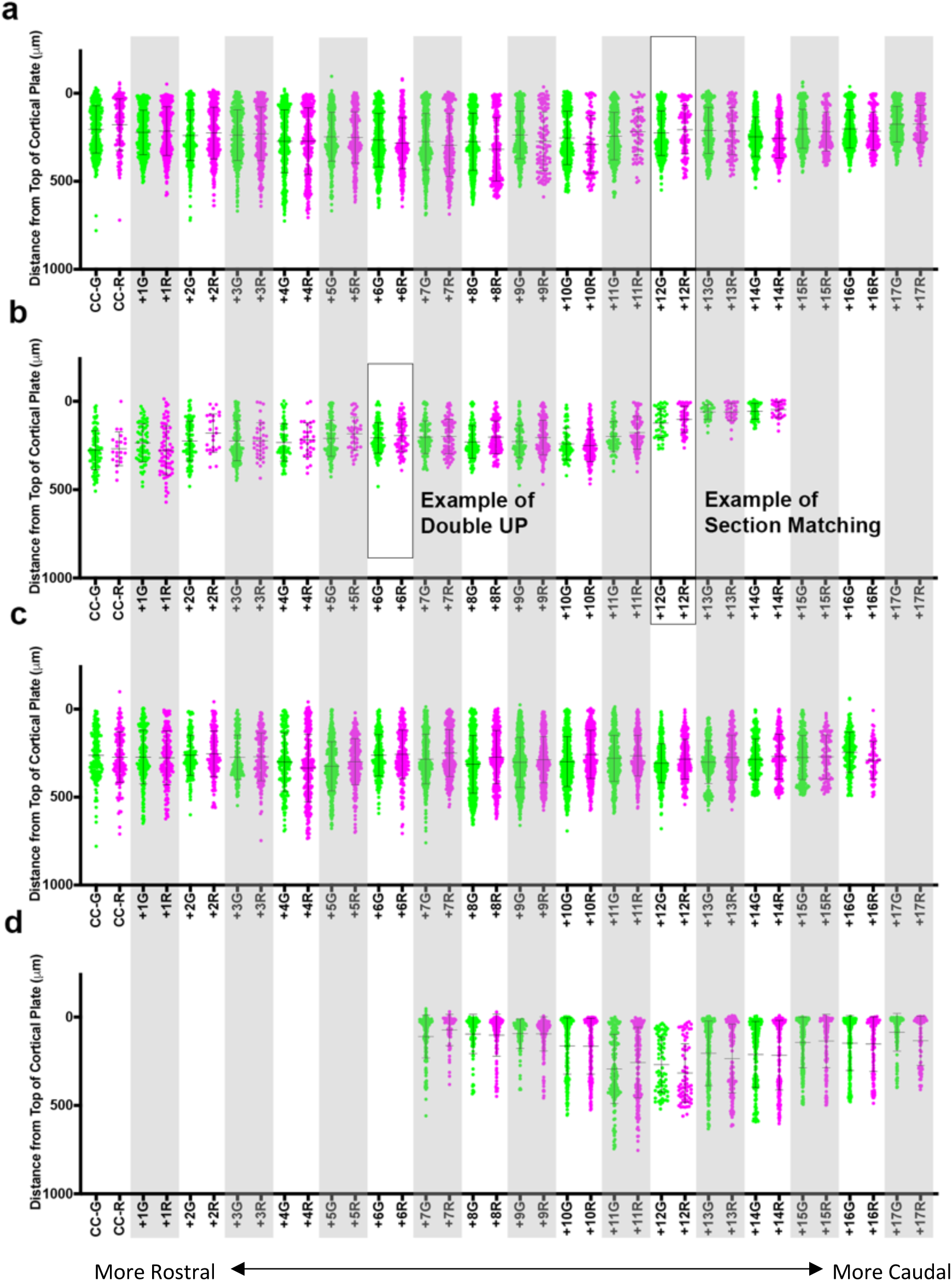
Double UP is superior to section matching. **(a-d)** Results for cell location of four different brains, all relative to the top of the cortical plate. Sections were collected and mounted sequentially and aligned across brains according to the first section which contained an uninterrupted corpus callosum (CC) (n=118-1026 cells per section, approximately half green, half red). This matched well with other common landmarks. For **Figure 1d**, Section Matching refers to all fluorescent cells (mNeon or mScarlet) from each section compared with each other perfectly matched sections, here shown as a boxed vertical column. “Double UP” refers to the mNeon or mScarlet cells within one section of one brain. Lines indicate mean and SD. G, R refer to all the green or all red cells within a section, respectively. +1, +2, +3 … refers to how many sections caudal of the CC the section being examined is located. Each section is 100µm.

**Supplementary Figure 3:**
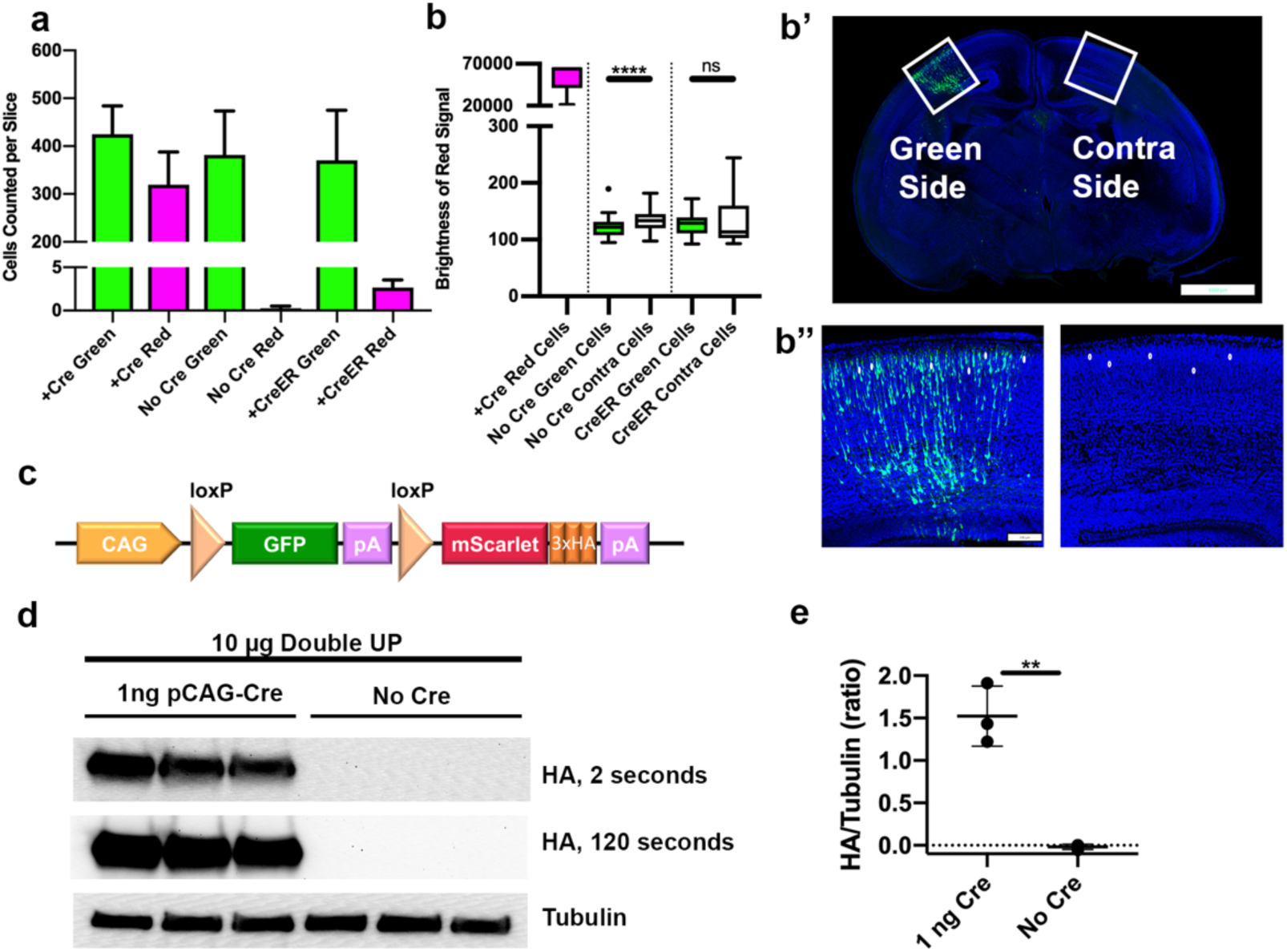
Double UP is not leaky. **(a)** Number of green and red cells of greater than 1000 units of brightness per slice in Double UP+Cre (3 slices, mean=425 green//320 red cells), Double UP-Cre (4 slices, mean=382 green /0.25 red cells) and Double UP +ERT2-Cre-ERT2 (3 slices, mean =370 green/3 red cells). Error bars indicate SEM. All images acquired with identical settings. **(b)** Quantification of the brightness in the red channel of red neurons from 90 neurons across three brains for: red neurons from Double UP+Cre, as well as green neurons from the electroporated (Green Side) and fluorescent negative neurons from the contralateral side (Contra Side) of brains having received Double UP without Cre and Double UP with CreER. Dot in “No Cre Green Cells” indicates an outlier, all other points contained within the whiskers (Tukey). Boxes are 25%/75%, median is shown. **** p<0.0001, unpaired two-tailed t-test. **(b’)** Representative image of coronal slice demonstrating green side and contra side. **(b”)** Representative image of green and contra sides used for analysis. **(c)** Schematic of Double UP-HA. **(d)** Western blots of protein from CAD cells transfected with 10µg of Double UP and the indicated amount of pCAG-Cre. Quantification done on HA exposed for 2 seconds, 120 second exposure only shown for reference. **(e)** Amount of HA protein detected in each condition. The average HA signal in three experiments was slightly less than background. ** p=0.0017, unpaired two-tailed t-test of three experimental replicates. Lines indicate mean and SD.

## METHODS

### Double UP Plasmids

We have generated multiple Double UP plasmids through standard cloning procedures. Full sequence information is available upon request, or through Addgene. Double UP mNeon-to-mScarlet is available through Allele Biotechnologies. Double UP superfolderGFP-to-mScarlet, and Double UP mClover3-to-mScarlet are available through Addgene (Addgene #120261 and #120262). We have also constructed and will deposit in Addgene: Double UP Halotag-to-mScarlet, Double UP RFP670-to-mScarlet, and Double UP iRFP720-to-mScarlet. This manuscript exclusively uses Double UP mNeon-to-mScarlet, but the others have been validated and work equally well, with different fluorescent properties.

pCAG-Cre and pCAG-ERT2-Cre-ERT2 were purchased from Addgene (Addgene #13375, #13777). For the internal control to be valid, it is essential that all plasmids contain the same promoter, and so these two are highly recommended for use in combination with Double UP.

### Mice

All mouse procedures were approved by the University of Wisconsin Committee on Animal Care and were in accordance with NIH guidelines. All mice were Swiss Webster strain and purchased from Taconic.

### In Utero Electroporation (IUE)

Timed matings were performed, and pregnant dams were used at embryonic day 14.5 (E14.5). Day E0.5 is the morning of sperm plug visualization. Plasmid DNA was mixed at a ratio of 75 copies of Double UP per copy of Cre, then mixed with Fast Green FCF (Sigma) to a final concentration of 2µg total plasmid/µl and loaded into pulled capillary needles. The dam was anesthetized with isoflurane, and a laparotomy was performed, exposing the embryos. The embryos were gently pulled out of the abdominal cavity. The capillary needles were inserted into the lateral ventricles, and approximately 0.25-0.5 µL DNA/Fast Green FCF was injected using a PicoSpritzer II (Parker Instrumentation). Electrical current was passed across the head, in five pulses of 40 volts each lasting 100ms on and 900ms off using a CUY21 Electroporator (Bex Co. LTD). After the last embryo was electroporated, the embryos were inserted back into the mother, and the laparotomy was sutured closed. Embryos were allowed to develop normally for 3 or 4 days, depending on experiment (E14.5+3 or E14.5+4).

### Tissue Collection

Embryos were again exposed via laparotomy on the mother after deep anesthesia with isoflurane. Embryos were removed from the uterus one by one, chest cavity was opened, a small incision was made on the right atrium and a needle was inserted into the left ventricle. Through this needle, the animal was perfused with approximately 1 mL of sterile saline and 3 mL of 4% paraformaldehyde (PFA) using an Instech perfusion pump, at the rate of approximately 1.25mL per minute. Following perfusion, heads were removed and left in PFA at 4°C overnight before dissection. After the last embryo was perfused, the dam was euthanized.

### Sectioning-Fixed Tissue

After 16 hours in PFA at 4°C, heads were transferred to PBS and brains were dissected. Brains were placed in 3% low melt agarose for 10 minutes at 42°C, then moved into 6% low melt agarose and allowed to set on ice. After the agarose hardened, the brains were sectioned on a Leica VT1000S vibratome at 100µm in PBS. Sections were stored for less than one week in PBS+0.2% sodium azide before being stained with 4′,6-diamidino-2-phenylindole (DAPI) and imaged.

### Section Preparation

DAPI was diluted to a final concentration of 2.4 nM in 0.4% Triton/PBS, and left on sections for one hour on a gently rotating platform at room temperature. After one hour, the sections were washed three times in PBS before being mounted in Aqua-Poly Mount (Polysciences). Slides were allowed to dry for at least one hour and then imaged within two days.

### Section Matching

When section matching was used, every 100µm section was collected and stored in sequentially. All sections were treated with DAPI and mounted. The first slice to contain an uninterrupted corpus callosum was identified, and set as slice 0. All sections containing fluorescent signal were imaged and analyzed. One brain contained fluorescent signal in sections located before the first slice to contain an uninterrupted corpus callosum, but these sections were not analyzed as there was no matching section in the other brains examined in this way.

### Imaging

Confocal imaging was performed on a Zeiss LSM 800. Images were acquired at 12 optical sections, each 1µm apart. 2×2 tiles were collected with a 20x/0.8NA Plan Apochromat objective, with 2x averaging. Unless otherwise specified, gain/laser power were altered between each image set to optimize image quality. Tiles were stitched together using the stitching tool in Zen, and resulting images were analyzed using the TRON Machine software described elsewhere in this manuscript.

### Immunoblotting

CAD cells, originally derived from a catecholaminergic neuronal tumor (Sigma) were cultured in DMEM/F12 (Gibco) with 8% FBS (Teak Serum) and 1% Pen-Strep (Gibco) and were transfected at 60% confluency with 10µg of Double UP and either no pCAG-Cre or 1ng pCAG-Cre using Lipofectamine 3000 (Invitrogen) following the manufacturer’s protocol. Cells were washed once with cold PBS before being lysed with 300µl NP-40 Lysis Buffer (Invitrogen) with Complete Mini (Roche) at 48 hours post-transfection. Lysate was spun at 21,000g for 10 minutes, and supernatants were flash-frozen and stored at −80°C until use. Samples were thawed and loaded onto a 4%-10% SDS-Page gel, then transferred to PVDF membrane (Millipore). Membranes were blocked in 5% milk in 0.1% TBS-T, incubated with primary antibody overnight at 4°C, and blotted with a HRP-conjugated secondary antibody for 1 hour. Antibodies used were mouse anti-HA (1:1000, sc-57592, Santa-Cruz), Mouse anti-Tubulin (1:10,000, T9026, Sigma) and goat-anti-mouse HRP (1:10,000, 115-035-174, Jackson). Protein bands were visualized using Pierce ECL Western blotting substrate (Thermo Scientific).

### TRacking Overlapping Neurons (TRON) Machine

The TRON machine was designed to perform reproducible, high throughput and largely automated calculation of cell body location and distance from one or more user defined regions of interest. It was specifically designed for progenitor cells and migrating neurons, or neurons that have recently completed migration. We anticipate it will work well for most cell types, excluding highly branched and differentiated neurons. Most steps are automated, manual steps will be indicated. The TRON machine largely utilizes ImageJ/Fiji tools. In short, images are processed via unsharp mask, gaussian blur, thresholding, erosion and watershedding to transform cells into reduced cell bodies. These cell bodies are then run through a modified 3D objects counter to determine which cells in a similar XY location in different slices are the same or distinct cells. The distance from each cell to a user defined region of interest is then calculated. A downloadable program, complete instructions for use and a sample Double UP image file can be found here: https://uwmadison.box.com/v/tron. Full code and associated instructions can be found here: **https://go.wisc.edu/ddy57r.**

### Statistics

The data shown in **Fig 1d** is shown in more complete detail in **Supp. Fig 2**. The representative scatter plots in **Fig 2e** are included in the graph and statistical calculations in **Fig 2f**. The same slices were examined for two different measurements in **Supp. Fig 3a, 3b**. All other measurements were from distinct samples. Data was tested for normality using the Kolmorgorov-Smirnov test of normality, if data was normal, two-tailed t-test was run, if data was not normal, the Kolmorgorov-Smirnov two-tailed t-test was run. Complete data is available upon request. All statistical tests were run in Prism 8 (Graph Pad).

## References

1. Saito, T. & Nakatsuji, N. Efficient gene transfer into the embryonic mouse brain using in vivo electroporation. Developmental biology 240, 237–246 (2001).

2. Tabata, H. & Nakajima, K. Efficient in utero gene transfer system to the developing mouse brain using electroporation: visualization of neuronal migration in the developing cortex. Neuroscience 103, 865–872 (2001).

3. Bayer, S. & Altman, J. Neocortical Development. (Raven Press., New York, New York; 1991).

4. Kootstra, N.A. & Verma, I.M. Gene therapy with viral vectors. Annu Rev Pharmacol Toxicol 43, 413–439 (2003).

5. Kilby, N.J., Snaith, M.R. & Murray, J.A. Site-specific recombinases: tools for genome engineering. Trends Genet 9, 413–421 (1993).

6. Shaner, N.C. et al. A bright monomeric green fluorescent protein derived from Branchiostoma lanceolatum. Nat Methods 10, 407–409 (2013).

7. Lanoix, J. & Acheson, N.H. A rabbit beta-globin polyadenylation signal directs efficient termination of transcription of polyomavirus DNA. EMBO J 7, 2515–2522 (1988).

8. Bindels, D.S. et al. mScarlet: a bright monomeric red fluorescent protein for cellular imaging. Nat Methods 14, 53–56 (2017).

9. Matsuda, T. & Cepko, C.L. Controlled expression of transgenes introduced by in vivo electroporation. Proc Natl Acad Sci U S A 104, 1027–1032 (2007).

10. Kawauchi, T., Chihama, K., Nabeshima, Y. & Hoshino, M. The in vivo roles of STEF/Tiam1, Rac1 and JNK in cortical neuronal migration. EMBO J 22, 4190–4201 (2003).

11. Konno, D., Yoshimura, S., Hori, K., Maruoka, H. & Sobue, K. Involvement of the phosphatidylinositol 3-kinase/rac1 and cdc42 pathways in radial migration of cortical neurons. J Biol Chem 280, 5082–5088 (2005).

